# MemSTATS: A Benchmark Set of Membrane Protein Symmetries and Pseudo-Symmetries

**DOI:** 10.1101/653295

**Authors:** Antoniya A. Aleksandrova, Edoardo Sarti, Lucy R. Forrest

**Author notes:** Correspondence to Lucy R. Forrest: Computational Structural Biology Section, National Institute of Neurological Disorders and Stroke, National Institutes of Health, 35 Convent Drive, Rm 3D991 MSC 3761, Bethesda, MD 20892-3761, USA. Biologie Computationnelle et Quantitative, CNRS - Sorbonne Université, Institut de Biologie Paris Seine, Case courrier 1540, 4 Place Jussieu, Paris 75005, France.

## Abstract

In membrane proteins, symmetry and pseudo-symmetry often have functional or evolutionary implications. However, available symmetry detection methods have not been tested systematically on this class of proteins due to the lack of an appropriate benchmark set. Here we present MemSTATS, a publicly-available benchmark set of both quaternary and internal symmetries in membrane protein structures described in terms of order, repeated elements, and orientation of the axis with respect to the membrane plane. Moreover, using MemSTATS, we compare the performance of four widely-used symmetry detection algorithms and highlight specific challenges and areas for improvement in the future.

---

Membrane proteins are encoded by around one third of a given genome [1–3] and play key roles in transmission of information and chemicals such as neurotransmitters into the cell. Available membrane protein structures have revealed an abundance of symmetry and pseudo-symmetry, which are observed not only in the formation of multi-subunit assemblies, but also in the repetition of internal structural elements. These symmetries are often intimately associated with the folding and function of membrane proteins and hence a systematic study of symmetry should provide a framework for a broader understanding of the mechanistic principles and evolutionary development of this important class of proteins.

Although sophisticated algorithms exist that detect repeated structural elements in proteins [4,5], there has been limited effort to investigate their applicability to membrane proteins. The performance of these algorithms for this class might differ for several reasons. First, the sequences of membrane proteins have diverged extensively [6,7], which can hinder the detection of internal symmetry. Second, the structures of some membrane proteins exhibit functional asymmetry [8], which might necessitate detection of multiple small, non-hierarchical symmetries to fully capture the relevant characteristics of the structure. However, investigations of the applicability of existing symmetry detection algorithms to membrane proteins are hampered by the lack of a reference benchmark that can be used to evaluate the performance of the detection methods. Previous attempts to create a benchmark set for protein symmetries have either excluded membrane proteins entirely [4] or have included very few representatives. For example, in a manually-curated dataset of 1007 symmetries in SCOP domains [5], only about 30 domains came from transmembrane proteins.

To understand how well-suited the available methods are for detecting membrane protein symmetries and to facilitate future investigations of the effect of the lipid bilayer on a protein’s predisposition for symmetry, we compiled a benchmark set of Membrane protein Structures And Their Symmetries (MemSTATS). We chose to describe membrane proteins in terms of complexes and their membrane-spanning chains rather than domains. While many reputable databases and benchmarks use domains as the fundamental unit [9,10], their definition for membrane proteins is often ambiguous [11]. Instead, we organize our data around protein chains, each being a single gene product [12], which naturally allows us to investigate both internal and quaternary symmetries in every entry of the dataset.

While compiling the MemSTATS set, we aimed to describe the diversity of membrane protein symmetries, while limiting the number of trivial cases such as a repeat composed of a single transmembrane helix. The resultant set comprises 87 α-helical and 23 β-barrel membrane proteins, each with distinct architecture either at the quaternary or internal level. For each symmetry, the following features are specified: symmetry order, axis orientation with respect to the membrane, approximate amino-acid range of the repeats, whether the symmetry is cyclic or open (as in helical or linear symmetries), whether the symmetry has been explicitly mentioned in the literature, and whether the repeats are interdigitating (https://doi.org/10.5281/zenodo.3234036). Unlike previous symmetry benchmarks, which included only the symmetry order, the more extensive descriptions of symmetry in MemSTATS make it possible to assess not only whether an algorithm detects a symmetry, but also how precisely the symmetry is characterized.

Using MemSTATS, we have assessed the performance of two well-established computational algorithms for detecting internal symmetry in protein structures, namely SymD [4] and CE-Symm [5,13], as well as two methods designed exclusively for quaternary symmetry detection, namely AnAnaS [14,15] and the BioJava module used for symmetry annotations of complexes in the protein data bank, PDB [16] (Fig. 1, 2, 3). We refer to the latter simply as BioJava.

**Figure 1.**
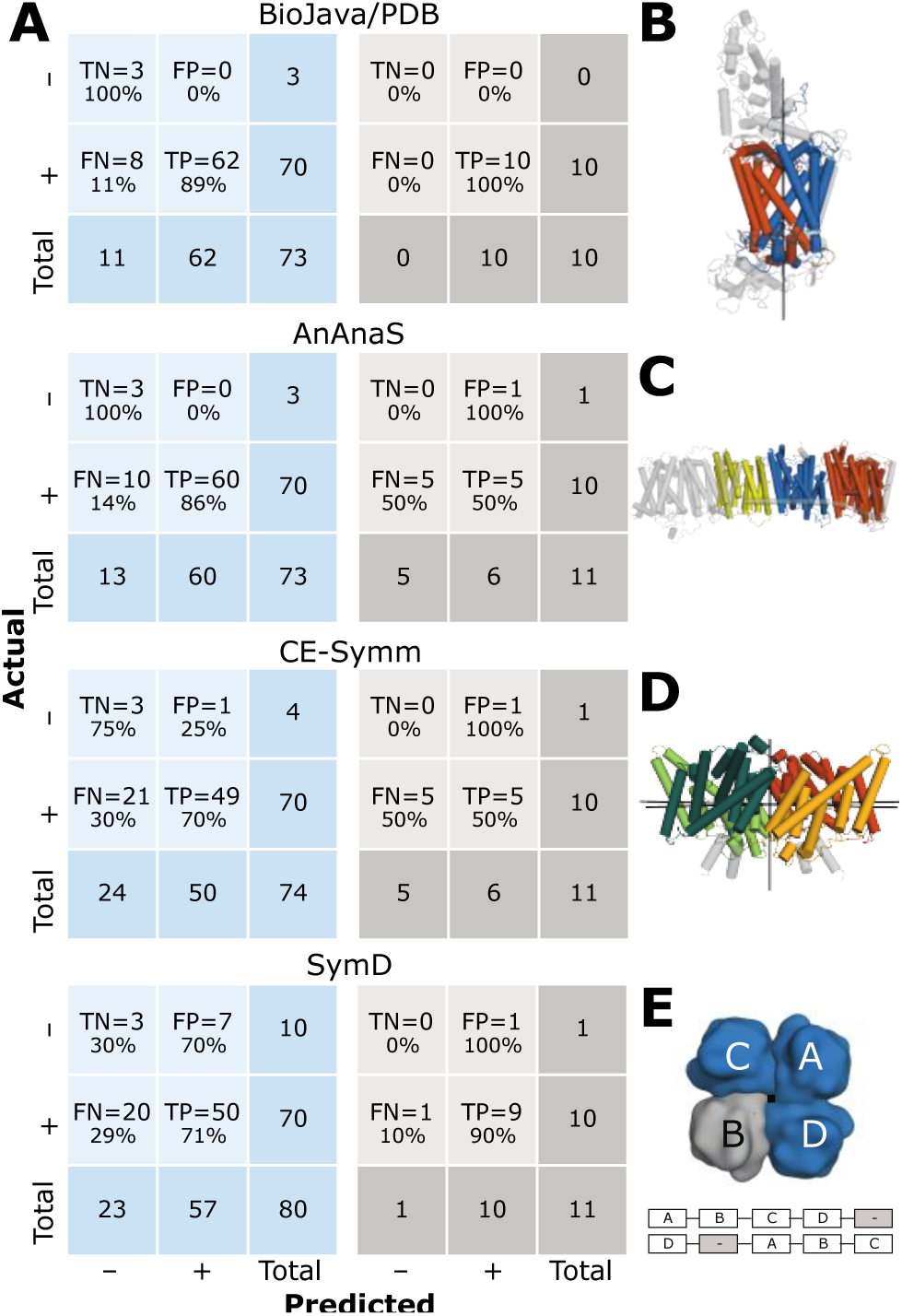
Quaternary symmetry detection in MemSTATS. **(A)** Confusion matrices summarizing the performance of each algorithm at detecting quaternary symmetries in the alpha-helical (blue) or beta-barrel (gray) transmembrane proteins in the MemSTATS dataset. SymD results were obtained using one of the two author-recommended thresholds: z-TM-score = 8. TN = true negatives, FP = false positives, FN = false negatives, TP = true positives. **(B)** Chain L (orange) and chain M (blue) in the bacterial photosynthetic reaction center (PDB 1prc) are structurally pseudo-symmetric (C_α_ RMSD ∼1.9 Å) around the shown axis (black) but have a very low sequence identity (∼22%), which prevents BioJava from detecting the pseudo-symmetry. **(C)** The membrane-embedded portion of Complex I (PDB 4hea), in which chains L (orange), M (blue) and H (yellow) are related by an open pseudo-symmetry not detectable by AnAnaS. **(D)** The dimer of the proton-chloride antiporter ClC (PDB 1ots) is C2-symmetric, but each of its protomers also contains a C2-pseudo-symmetry. CE-Symm detects all of these symmetries but only insofar as they are hierarchically related to the smallest repeated subunits (shown in different colors). Because the internal pseudo-symmetries do not encompass all the residues in the protomers, CE-Symm does not report the full extent of the quaternary symmetry: the undetected portions are in gray. **(E)** Internal symmetry detection programs CE-Symm and SymD merge all chains in a complex according to the order in which they appear in the submitted coordinate file. If this sequence does not correspond to the structural proximity of the chains, the ability of these algorithms to detect even a perfect C4 symmetry, like the one found in a potassium KcsA channel (PDB 1k4c), is hampered. For instance, in the example shown, SymD might be able to detect that the blue-colored subunits are related by a C4 symmetry transformation but would not find a match for the gray subunit.

**Figure 2.**
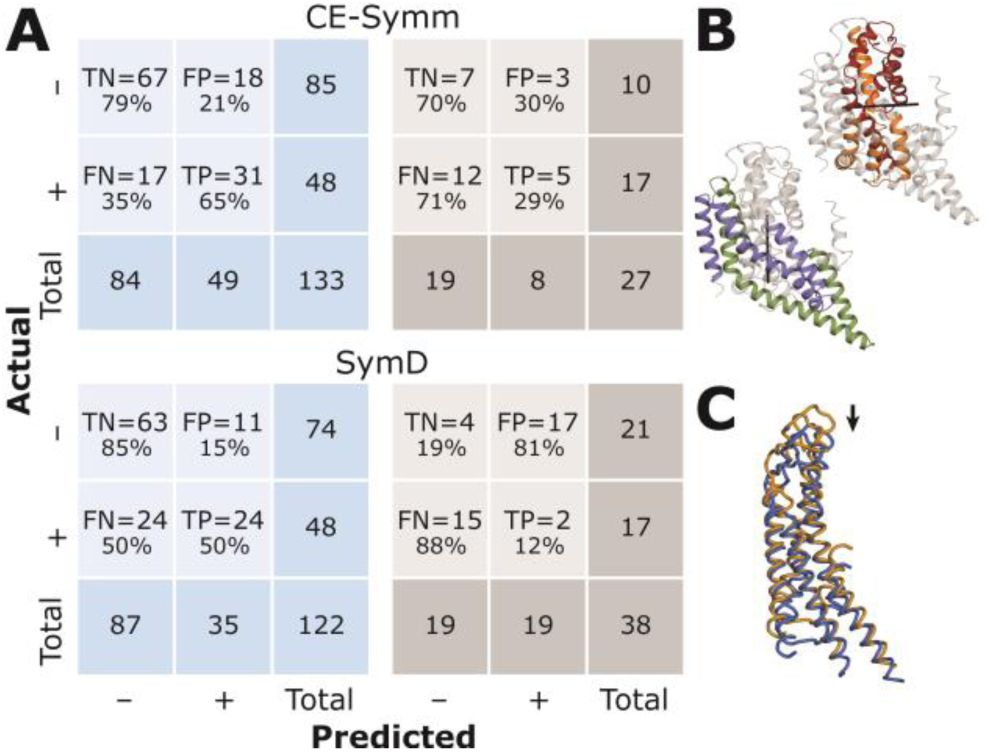
Internal symmetry detection in MemSTATS. **(A)** Confusion matrices summarizing the performance of each algorithm at detecting internal symmetries in the alpha-helical (blue) and beta-barrel (gray) transmembrane proteins in the MemSTATS dataset. SymD results were obtained using one of the two author-recommended thresholds: z-TM-score = 8. For beta-barrel proteins, structural internal symmetry has poorly-understood functional or evolutionary meaning and, hence, these cases were included only for completeness in the benchmark and are based on number of strands. TN = true negatives, FP = false positives, FN = false negatives, TP = true positives. **(B)** Symmetries with independent axes in the same structure shown for chain A of the glutamate transporter homolog Glt_Ph_ (PDB 2nwx). None of the discussed detection algorithms can simultaneously detect both axes. **(C)** Example of so-called slip symmetry: a trivial superposition of chain A from the complex of a connexin 26 gap junction channel (PDB 2zw3) (blue) onto its copy that has been shifted by 20 residues (orange) can receive a high structural alignment score. Such slip symmetries are a frequent source of false positives for SymD.

**Figure 3.**
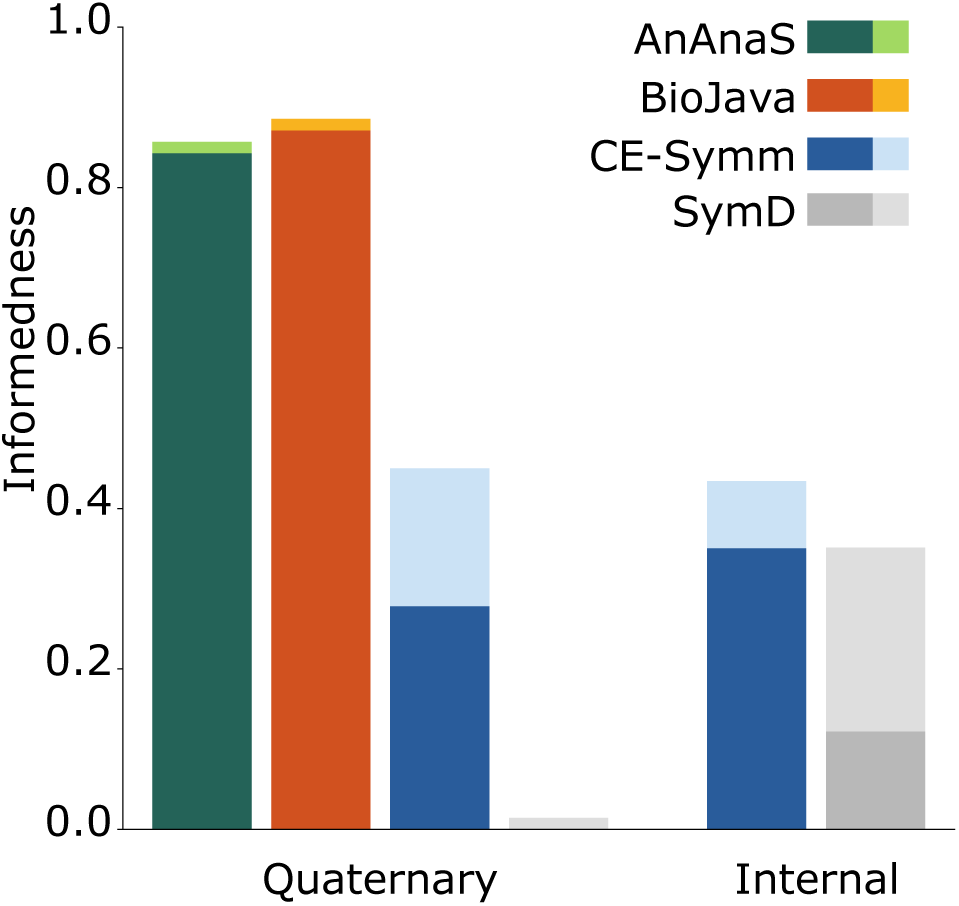
Comparison of the performance of symmetry detection algorithms on the symmetries in the alpha-helical transmembrane proteins in the MemSTATS dataset. Informedness is used as a metric of accuracy and is calculated as True Positive Rate + True Negative Rate – 1. Informedness ranges from 0 to 1, with 1 representing perfect performance of the algorithm and 0 indicating an inability to distinguish between true and false predictions. Colors indicate the four methods: green for AnAnaS, orange for BioJava, blue for CE-Symm, and gray for SymD computed with z-TM-score cutoff of 8. The lighter shade indicates results for which the symmetric order was detected correctly but only a fraction of the expected symmetry-related residues were recognized (with an allowed error of 20 consecutive residues). For SymD, changing the z-TM-score cutoff to 10 increases the informedness for quaternary symmetries from 0.01 to 0.03 and decreases it for internal symmetries from 0.35 to 0.27.

The two quaternary-symmetry detection methods, BioJava and AnAnaS, differ significantly in their approach for detecting the point group and symmetry axes for a complex. However, both methods strive to optimize the speed of repeat detection by taking advantage of the fact that in quaternary symmetry there is often a correspondence between repeat and chain boundaries. Therefore, to identify symmetry-related subunits, the two algorithms rely on sequence alignment and structural superposition of each pair of chains. While this strategy makes them suitable to efficiently scan a large number of protein complexes, it also means that, by design, AnAnaS and BioJava cannot detect symmetries within a protein chain.

On the other hand, SymD and CE-Symm both specifically target internal symmetries. To identify structurally similar regions of the protein chain, these two algorithms align the protein structure to itself. For example, SymD aligns two copies of the same protein, circularly permutes the second copy, one residue at a time, and scores the resulting structural superpositions. This approach allows SymD to deduce the order and axis of symmetry, as well as the number of residues in the protein that are symmetry-related, but not the boundaries of the repeats. In contrast, CE-Symm uses dynamic programming to align small fragments of the protein and then applies a series of thresholds (e.g. of fragment size) to identify a global structural self-alignment from which both symmetry axes and repeat boundaries can be extracted. Designed with internal symmetry in mind, both SymD and CE-Symm implicitly assume that connected, repeated elements tend to be not only sequentially but also spatially proximate. This assumption, however, can make it challenging to use SymD or CE-Symm for detecting the order of quaternary symmetry in complexes with more than three symmetry-related repeats, since all possible arrangements of the repeats need to be taken into account. Therefore, SymD and CE-Symm, which process complexes as a single polypeptide comprised of the chains concatenated in the order of their appearance in the PDB file, struggle to detect symmetry in large complexes in which the names of the chains do not correspond to their spatial proximity (e.g., PDB identifier PDB: 1K4C; Fig. 1E). Note that AnAnaS and BioJava can both correctly characterize such cases.

While MemSTATS allows for the four algorithms to be benchmarked against each other, it should be noted that the set is relatively small due to the current availability of structures and the emphasis on avoiding redundancy in the types of challenges presented. Therefore, overall metrics for scoring should be carefully selected to take into account the size and imbalance in the dataset. With this in mind, we summarized the key results for each method using confusion matrices (Fig. 1A, 2A) and also compared the probability that each algorithm produces informed predictions relative to chance (Fig. 3). These results indicate that while most methods detected more symmetries than they miss, a substantial fraction of symmetries were not detected, especially for cases of internal symmetry. In addition, it is clear that identifying the boundaries of internal repeats is a non-trivial problem (Fig. 3). To better understand the specific situations in which each method performs well or poorly, we used MemSTATS to examine individual examples in the dataset.

AnAnaS, for example, due to its analytic framework, is limited to global cyclic, dihedral, or cubic symmetries, and therefore did not detect either open symmetries, such as helical or linear symmetries (e.g., PDB: 4HEA; Fig. 1B), or local symmetries, in which only a subset of chains in a complex are related (e.g., PDB: 3WMM, PDB: 2R6G, PDB: 4Y28). All other methods can, in principle, detect such local and open symmetries. BioJava, on the other hand, failed for cases where the symmetry-related chains scored below the assigned threshold of 40% sequence identity (e.g., PDB: 1PRC, PDB: 1ZOY, PDB: 2R6G, PDB: 4Y28, PDB: 4HEA; Fig. 1C). Such cases were typically correctly identified by the internal symmetry algorithms, as long as the repeats constituted a large fraction of the complex (e.g., PDB: 2R6G but not PDB: 4HEA).

The ability to detect open symmetry makes internal symmetry algorithms vulnerable to obtaining a well-scoring self-alignment by just a slight translation of the structure. This so-called “slip” symmetry matches most secondary structure elements from the transformed structure to themselves and, hence, proteins with an abundance of straight parallel alpha-helices or beta-strands, for example, were especially prone to elicit this behavior (Fig. 2C). While the various thresholds assigned in SymD and CE-Symm attempt to mitigate the detection of false positives due to slip symmetry, the problem persisted for SymD in a number of cases (e.g., PDB: 4HEA, PDB: 2ZW3_A).

Cases with symmetry mismatch proved to be inscrutable to all the abovementioned algorithms. Such cases included the structures of the AMPA (PDB: 4U1W) or NMDA (PDB: 4PE5) receptors, which besides the readily-detectable global two-fold symmetry of the complex, also exhibit two unrelated and distinct pseudo-symmetries relating the chain fragments in the membrane-embedded and extracellular components of the complex (Fig.2 in [17]). These symmetries presented a challenge for AnAnaS and BioJava because neither method considers repeats formed by a fraction of a chain. Among the internal-symmetry detection methods, SymD is inherently unable to detect more than one symmetry in a structure, while CE-Symm can only detect multiple symmetries simultaneously if those symmetries are hierarchical, i.e., they relate sub-regions of the same symmetric repeats. These limitations of the internal-symmetry detection algorithms also prevented them from identifying two distinct symmetries within a single protein chain that cannot be related to one another, such as those found in the protomer of a glutamate transporter homolog (Glt_Ph_, PDB: 2NWX_A; Fig. 2B).

One of the main challenges for internal symmetry algorithms is detecting the full extent of repeats. First, since there is often much more sequence and structure divergence between repeats in internal symmetry than in quaternary symmetries, setting appropriate thresholds for determining whether a protein structure is symmetric and which residues are involved can be difficult. This issue was apparent for protein structures in which a pronounced structural divergence between repeats is of functional significance, such as in vcINDY (PDB: 4F35_C) [18,19]. Second, CE-Symm, due to its focus on detecting multiple, hierarchically-related symmetries within a structure, might only identify some of the residues of the repeats related by a given symmetry axis because the remainder do not satisfy the criteria of a second symmetry axis. Such an outcome was observed not only within protein chains (e.g. PDB: 3NCY_A), but also for complexes that exhibit both quaternary and internal symmetry (e.g. PDB: 1OTS; Fig. 1D).

In conclusion, the MemSTATS data set provides a solid foundation for evaluating both the overall performance of symmetry detection algorithms and the specific challenges in membrane protein structures that each method can address. However, the limitations of the dataset, including its relatively small size, should be kept in mind when interpreting the statistical significance of results obtained thereon. For example, at the quaternary level, more than one complex within the dataset has similar architecture (e.g., there are multiple examples with C4 quaternary symmetry), and therefore the specific issues that this architecture presents will be upweighted when judging the overall performance of an algorithm. Furthermore, only a few multi-chain proteins have no quaternary symmetry, while few beta-barrel proteins have no internal symmetry; this imbalance means that measures such as accuracy should be avoided or used with care. We also note that the determination of internal structural symmetry in beta-barrels is highly ambiguous, because it is not fully apparent how the symmetry might relate to function; these cases were included in the dataset mainly for completeness. The abovementioned issues are unlikely to be solved by the increasing number of membrane protein structures since it is likely that they are intrinsic to the types of symmetries that can be encountered within the membrane environment. Nevertheless, it is likely that the discovery of new architectures will pose novel and interesting challenges to symmetry detection algorithms.

## Acknowledgements

This research was supported by the Division of Intramural Research of the NIH, National Institute of Neurological Disorders and Stroke. We thank the LOBOS administrative team of the National Heart, Lung and Blood Institute for computational support.

## Appendix A. Supplementary data

The current release of the MemSTATS data set is available online at: https://doi.org/10.5281/zenodo.3234036. The dynamic version of the data set, as well as the Python code used to benchmark the results of the symmetry detection methods, is available online at: https://github.com/AntoniyaAleksandrova/MemSTATS_benchmark. The raw outputs of the benchmarked symmetry-detection algorithms can be found at https://doi.org/10.5281/zenodo.3228540.

## Abbreviations

MemSTATS: Membrane protein Structures And Their Symmetries

